# Explainable Prototype Booster: Enhancing Latent Representations of Foundation Models for Gene Expression Prediction

**DOI:** 10.64898/2026.04.27.720478

**Authors:** Chaoyi Li, Quan Nguyen

**Author notes:** Contributing authors.

## Abstract

Spatial transcriptomics (ST) is a cutting-edge technology that measures gene expression while preserving spatial context and generating pathology-grade tissue images. Although ST has enabled numerous discoveries and demonstrated a huge application potential in pathological diagnosis and prognosis, the technology remains time-consuming and costly. The ability to predict gene markers of cancer from histological H&E-stained tissue images can overcome these technological barriers to open new horizons for precision and personalised pathology. Recently, foundation models have demonstrated improvements in generating general-purpose embeddings of H&E-images. However, these improved representations are not optimized for gene expression prediction and lack task-specific adaptability. To address this limitation, we propose the Explainable Prototype Booster (EP-Booster), which incorporates biological prior knowledge to guide the construction and training of learnable prototypes for embedding refinement, thereby improving gene expression prediction. Importantly, model predictions are inherently interpretable through pathway-level attributions associated with the prototypes. Extensive experiments across multiple datasets, cancer types, and spatial transcriptomics platforms demonstrate that EP-Booster consistently outperforms existing methods. Moreover, EP-Booster can be integrated with diverse foundation models to enhance task-specific representations, thereby improving predictive performance and biological interpretability in clinically relevant applications, including cancer biomarker prediction, survival analysis, and drug response prediction.

## Introduction

Spatial transcriptomics (ST) technologies, such as Visium, add spatially-resolved molecular information to H&E-stained images [1]. The combination of morphological features and molecular profiles reveals cellular heterogeneity across tissue regions, a breakthrough recognized as the “Method of the Year” [2], highlighting its significant potential in both basic and translational research. However, current ST protocols are costly and labor-intensive, posing challenges for large-scale data generation and for clinical deployment. To address the challenges, several modelling approaches have been developed to predict gene expression profiles directly from H&E-stained images using deep learning techniques. For instance, ST-Net [3] used a DenseNet-121 backbone pre-trained on ImageNet to extract image features, followed by a fully connected layer to predict gene expression. HisToGene [4] incorporated spatial context by linking patch features to their tissue locations, while TRIPLEX [5] further improved performance through multi-scale feature integration. However, because these models rely solely on extracting image information, their embeddings often fail to align well with gene expression. To mitigate this gap, BLEEP [6] employed contrastive learning to embed both modalities into a shared space and retrieve expression profiles via nearest-neighbor search.

Recently, pathology foundation models (e.g., UNI [7] and CONCH [8]) were leveraged to extract image embeddings from H&E-stained slides for predicting gene expression. This line of work benefits from the strong representational power of foundation models but still yields suboptimal results. Foundation models are primarily trained with self-supervised learning on large-scale pathology image collections, enabling their embeddings to capture broad morphological patterns. However, these embeddings often contain redundant information and are not explicitly optimized for downstream tasks such as gene expression prediction. To mitigate this limitation, Stem [9] employed joint distribution modeling to align gene expression and image embeddings in a shared latent space, but its effectiveness depends heavily on large training datasets. As an alternative, dimensionality reduction techniques such as Principal Component Analysis (PCA) reduce high-dimensional data by projecting it onto directions of maximum variance, thereby preserving the most informative structures. Experiments on the HEST benchmark [10] show that applying PCA to image embeddings from pathology foundation models improves gene expression prediction, suggesting that the original embeddings may contain task-irrelevant components that hinder prediction performance. However, by emphasizing high-variance directions, PCA may suppress biologically meaningful features with lower variance, potentially reducing interpretability. Therefore, developing an effective strategy to directly reconstruct these embeddings by incorporating task-specific biological information has great potential to improve predictive performance.

On the other hand, prototype learning has gained significant attention in computational pathology and has become a widely adopted approach for H&E image analysis. A prototype refers to a representative instance that captures the shares characteristics of data points belonging to the same type or category, and is often used to encode semantic or conceptual information. For example, AttnMISL [11] and H2T [11] perform k-means clustering on patch embeddings and use the resulting cluster centroids as prototypes. PANTHER [12] employs a Gaussian mixture model (GMM) to summarize patches from whole-slide images (WSIs) into a set of compact morphological prototypes. MMP [13] extended PANTHER by incorporating pathway-level prototypes and integrating them with morphological prototypes for downstream tasks. In contrast to these approaches, we propose to construct learnable prototypes directly guided by biological information. These prototypes are used to reconstruct image embeddings from foundation models, thereby enhancing their utility for gene expression prediction.

Motivated by these observations, we propose a lightweight module called Explainable Prototype Booster (EP-Booster), which incorporates biological prior knowledge to guide the construction and learning of task-specific prototypes, thereby refining image embeddings from foundation models. By reconstructing the embeddings through EP-Booster, task-irrelevant information is removed, while biologically meaningful semantic features are preserved, leading to improved gene expression prediction. At the core of EP-Booster is a prototype-guided cross-attention module that facilitates interaction between image embeddings and learnable prototypes. In addition, the joint supervised reconstruction–regression objective in EP-Booster refines the image embeddings by preserving essential visual information while simultaneously injecting biologically meaningful signals derived from selected gene expression profiles. Furthermore, we introduce a prototype initialization strategy that includes both random initialization and gene program-guided initialization, offering flexibility to integrate either solely data-driven signals or biologically informed priors during training, thereby establishing a strong foundation for learning task-specific prototypes tailored to gene expression prediction. For gene program–based initialization, gene programs are constructed by integrating spatially variable genes (SVGs) with curated biological pathways. SVGs capture spatial heterogeneity in gene expression, while pathways encode coordinated biological functions, resulting in prototypes that are biologically meaningful and well aligned with image embeddings. Building on gene program–guided initialization and prototype-guided cross-attention, we further develop a pathway-level interpretability method that maps model predictions back to biologically relevant pathways. EP-Booster does not modify or adapt any weights of the foundation model. Instead, EP-Booster operates solely on top of frozen image embeddings, learning external parameters that refine the embedding space without altering the backbone. Furthermore, EP-Booster is designed to remain effective under extremely limited sample regimes (such as spatial transcriptomics), where updating backbone parameters is impractical [12].

The main contributions of this work can be summarized as follows:

- EP-Booster is a new approach to spatial digital pathology with explainable prototype-based cross-attention module that directly reconstructs image embeddings from foundation models by incorporating biological prior knowledge;
- We introduce an alignment solution using a joint reconstruction-regression optimization strategy to enable EP-Booster to refine image embeddings that preserve essential visual information while injecting biologically meaningful signals to improve gene expression prediction;
- We propose a prototype initialization strategy, where gene program-based initialization imparts biological semantics to prototypes and improves their interaction with image embeddings;
- We propose a pathway-level interpretability method that links model predictions to relevant biological pathways;
- Our extensive experiments demonstrate that EP-Booster achieves superior performance and strong generalizability across diverse datasets and foundation models, while also proving effective for gene expression signature prediction;
- EP-Booster can predict clinically relevant outcomes, including survival analysis and drug responses, robustly validated across external datasets.

## Results

### Problem Formulation for Gene Expression Prediction

Given a set of H&E image patches *I*_*N*_ ∈ ℝ^*H*×*W* ×3^ and the corresponding gene expression profiles *G* ℝ^*N* ×*C*^, where *H* × *W* denotes the spatial resolution of each patch, *C* is the number of genes, and *N* is the number of patches. We further define 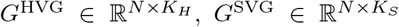 and 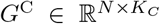 as subsets of *G* corresponding to highly variable genes (HVGs), spatially variable genes (SVGs) and clinically relevant markers, respectively, where *K*_*H*_, *K*_*S*_ and *K*_*C*_ denote the number of selected genes in each category.

For gene expression prediction, existing pipelines generally fall into two categories. The first directly applies an MLP on image embeddings 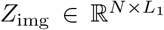 extracted by a pathology foundation model or a custom image encoder. The second also relies on foundation model embeddings but employs a regression model (e.g., ridge regression) to predict gene expression *G*. As regression models offer closed-form solutions with clearer interpretability compared to MLPs, our approach adopts the second paradigm. Building upon this framework, we introduce EP-Booster, a lightweight and explainable module that refines *Z*_img_ into a latent embedding 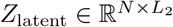 through a prototype-based representation learning approach. Specifically, learnable prototypes initialized from gene programs interact with image embeddings to inject biologically meaningful signals, producing task-aligned representations for gene expression prediction.

### EP-Booster Learnable Prototype Architecture for Gene Prediction

An overview of the EP-Booster pipeline is illustrated in Figure 1. During training, the model takes as input a triplet {*I*_*N*_, *G*^HVG^, *G*^SVG^}. First, a foundation model extracts image embeddings *Z*_img_ from the H&E image patches *I*_*N*_. Next, EP-Booster introduces a set of learnable prototypes 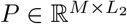, which are initialized using a prototype initialization strategy (either randomly or informed by gene programs derived from *G*^SVG^). EP-Booster then refines the image embeddings by projecting *Z*_img_ into a biologically informed latent space *Z*_latent_ through interactions between image features and the learnable prototypes. The model is optimized using highly variable genes *G*^HVG^ and *Z*_img_ as supervision signals. In this design, SVGs and HVGs serve complementary roles. SVGs are used for prototype initialization because they capture spatial heterogeneity within tissues, providing biologically meaningful priors that align well with morphological patterns in histology images. In contrast, HVGs are used for supervision, as their large expression variability provides stronger optimization signals, enabling more stable gradient updates and more effective learning of task-relevant representations. As a result, *Z*_latent_ encodes not only visual patterns but also biologically meaningful information aligned with gene expression profiles. After training, the refined latent embeddings *Z*_latent_ are paired with their corresponding target gene expression profiles (*e*.*g. G*^C^) to train a regression model.

**Fig. 1.**
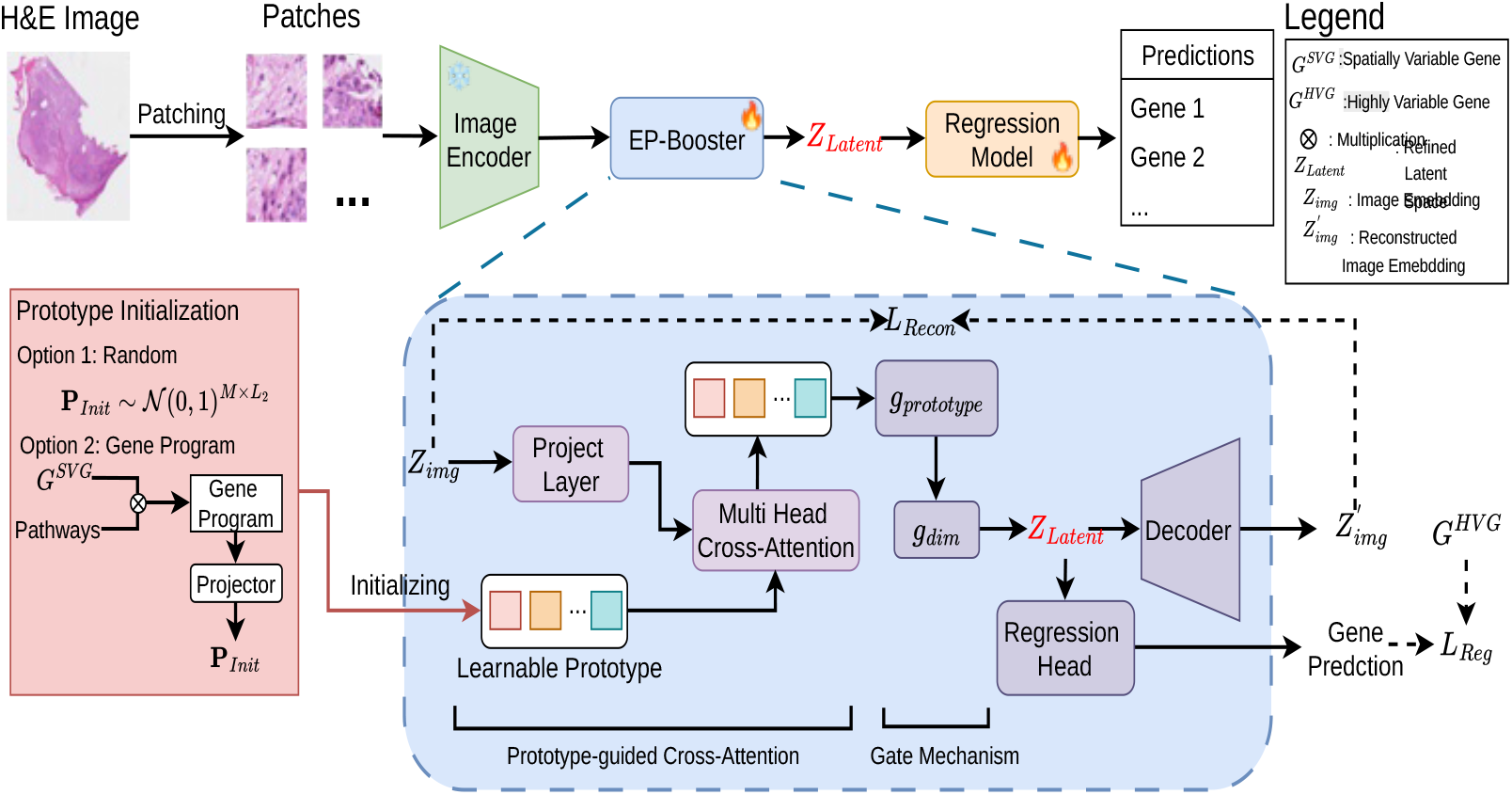
The EP-Booster Algorithm. The H&E image is split into patches, and each patch is encoded into *Z*_img_ using a frozen pathology foundation model. Learnable prototypes are initialized through a prototype initialization strategy and combined with *Z*_img_ via cross-attention and gating to produce refined embeddings *Z*_Latent_. These embeddings are optimized with reconstruction and regression losses guided by *Z*_img_ and *G*^HVG^, and subsequently used to train a regression model for gene expression prediction. The pseudo-code for **EP-Booster** is provided in the Appendix 1.

At inference time, only H&E image patches are required. These patches are processed by the foundation model and the trained EP-Booster to obtain *Z*_latent_, which is subsequently fed into the regression model to predict target gene expression. However, a central challenge in this framework lies in reconstructing *Z*_img_ using biological priors. Inspired by Q-Former [14], which extracts task-relevant visual features for text generation via learnable queries, we propose EP-Booster as a biologically driven refinement module tailored for this task. EP-Booster comprises four key components: a prototype-guided cross-attention module, prototype and latent gating mechanisms, a decoder, and a regression head. Details are provided in Section Methods.

### EP-Booster Outperforms State-of-the-Art Models

We compare EP-Booster with existing methods, including ST-Net, HisToGene, TRIPLEX, STEM, and BLEEP, across two types of gene expression prediction tasks: (i) prediction of the top 50 highly variable genes (HVGs), and (ii) prediction of genes with both high mean and high variability (HMHVGs).

As shown in Figure 2a and b, EP-Booster demonstrates consistent performance gains over existing methods on the HVG prediction task across both datasets, irrespective of the underlying foundation model (Figures S1, S2). Consistently, the superior performance was also observed by using mean square errors (Table S1). This demonstrates the robustness of our approach across different cancer types on the Xenium data. In addition, prototype initialization guided by gene programs (specifically those derived from SVGs) consistently yields better performance than random initialization as described in Table 1. This indicates that spatially informed gene programs more effectively capture underlying spatial patterns, thereby facilitating more meaningful embedding reconstruction.

**Table 1.** Generalization performance of the proposed method across various foundation models on the SKCM and LUAD datasets from the HEST benchmark [10]. Reported results are the mean values obtained from cross-validation. The ∗ denotes results reported by [10]. R is the Ridge regression model.

| Method | SKCM(HVGs) PCC ↑ | | | | LUAD(HVGs) PCC ↑ | | | |
| --- | --- | --- | --- | --- | --- | --- | --- | --- |
|  | R* | R + PCA* | R + Ours_R | R + Ours_GP | R* | R + PCA* | R + Ours_R | R + Ours_GP |
| ResNet 50 | 0.4117 | 0.4822 | 0.5139 | **0.5297** | 0.4001 | 0.4917 | 0.5406 | **0.5459** |
| CTransPath | 0.4038 | 0.5196 | 0.5512 | **0.5572** | 0.4026 | 0.4985 | 0.5745 | **0.5749** |
| Phikon | 0.3684 | 0.5355 | 0.5557 | **0.5729** | 0.4224 | 0.5468 | 0.5868 | **0.5973** |
| CONCH | 0.5079 | 0.5791 | **0.5811** | 0.5642 | 0.4957 | 0.5312 | 0.5566 | **0.569** |
| GigaPath | 0.2231 | 0.5538 | 0.5704 | **0.5706** | 0.3144 | 0.5399 | 0.5594 | **0.562** |
| UNI | 0.3433 | 0.6254 | **0.6337** | 0.6239 | 0.3714 | 0.5511 | 0.5565 | **0.5795** |
| Virchow | 0.3389 | 0.6088 | 0.6464 | **0.6573** | 0.2897 | 0.5459 | 0.6067 | **0.612** |
| Virchow2 | 0.303 | 0.6174 | 0.6482 | **0.6522** | 0.3017 | 0.5605 | 0.5837 | **0.60** |
| H-Optimus-0 | 0.2778 | 0.6432 | 0.6571 | **0.6632** | 0.3143 | 0.5582 | 0.5875 | **0.5907** |
| Average | 0.3531 | 0.5739 | 0.5953 | **0.599** | 0.368 | 0.536 | 0.5725 | **0.5813** |

**Fig. 2.**
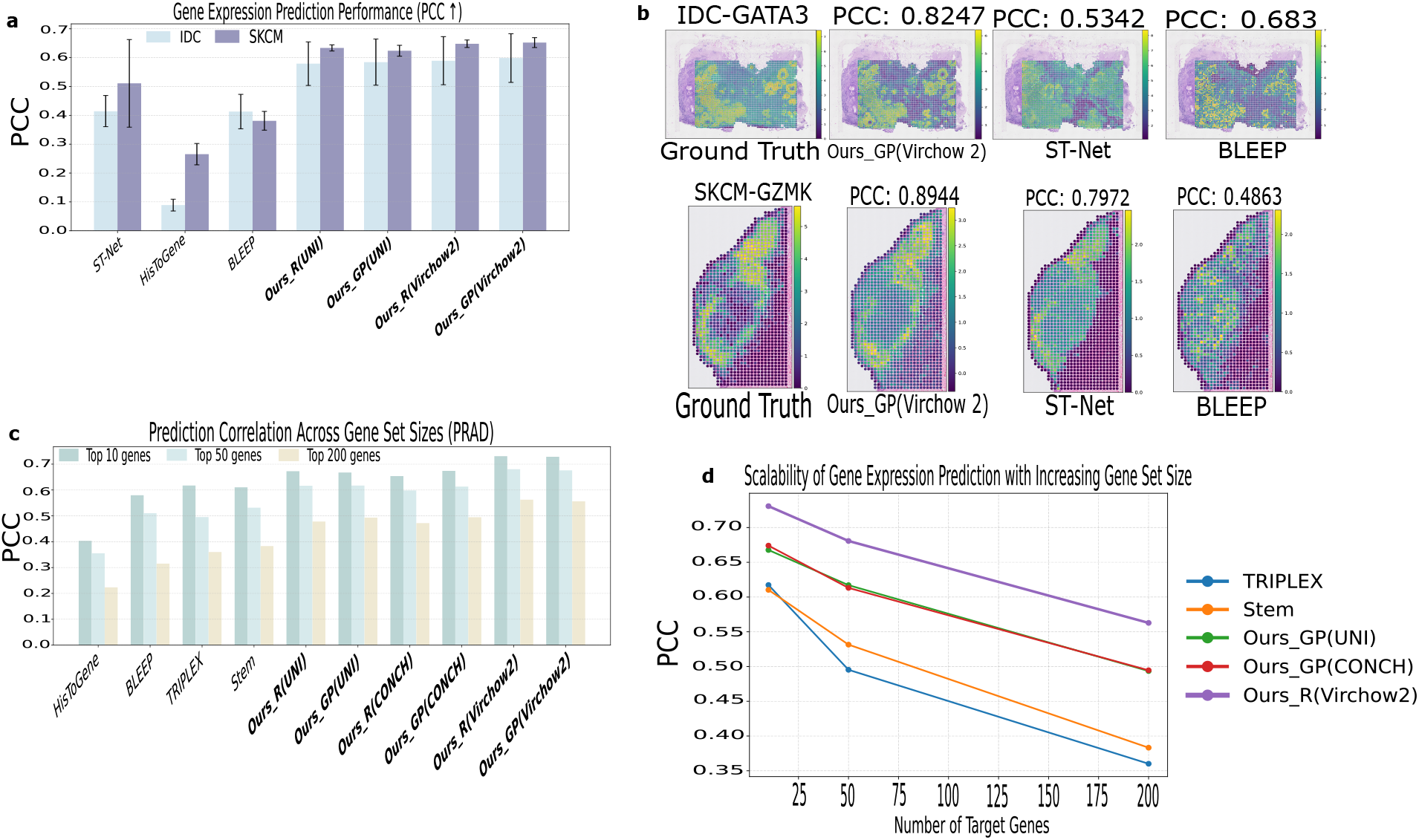
EP-Booster Performance. **a**, Compare EP-Booster with state-of-the-art methods on the IDC and SKCM datasets from the HEST benchmark [10] for predicting top 50 HVGs, evaluated using PCC. Reported results are presented as the mean and standard deviation across cross-validation folds. Ours R denotes EP-Booster with random prototype initialization, while Ours GP denotes EP-Booster with gene program–guided initialization. **b**, Visualization of predicted expression levels for HVGs (*e*.*g*.GATA3 and GZMK) produced by different methods on the IDC and SKCM datasets. **c**, Comparison of EP-Booster with state-of-the-art methods on the PRAD dataset from the HEST benchmark [10] for predicting HMHVGs. Reported results correspond to mean performance values obtained from cross-validation across varying gene set sizes. Results of current methods as reported in [9]. **d**, Scalability analysis of gene expression prediction as the gene set size increases, comparing different models.

To further evaluate the generalization capability of the proposed method across different ST platforms and varying gene set sizes, we conduct an additional experiment on a Visium PRAD dataset for predicting HMHVGs. Figure 2c shows that EP-Booster significantly outperforms other methods in all settings, indicating its robustness across both platforms and prediction scales. Moreover, consistent with findings in Figure 2a, prototype initialization based on gene programs generally yields better performance than random initialization. However, an exception is observed when using Virchow 2, where random initialization is slightly better than gene program-based initialization. We assume the reason could be the lower spatial resolution of the Visium with a mixture of cell types per spatial spot, where prior information from gene programs may restrict the representation ability of prototypes, limiting their ability to adapt to the complex latent space from Virchow 2. As shown in Figure 2d, although predictive performance decreases for all methods as the number of target genes increases, the rate of degradation varies substantially across models. Current methods, including TRIPLEX and STEM, exhibit a pronounced performance drop when the target gene set reaches 200 genes. In contrast, EP-Booster demonstrates markedly improved scalability, maintaining relatively high correlation levels even under the most challenging conditions.

### Generalization for Foundation Models

To evaluate the adaptability of EP-Booster to different foundation models, we compared it with the PCA reduction on the SKCM and LUAD datasets. As shown in Table 1, applying PCA substantially improves the performance of foundation models, yielding average PCC gains of 62.5% and 45.7% on the two tasks, respectively, compared to using the foundation models alone. This improvement is likely since raw image embeddings from foundation models contain a considerable amount of task-irrelevant or redundant information, which hinders effective learning by the regression model. Next, we notice that EP-Booster (with both random and gene program-based initialization) consistently outperforms PCA. For example, our method with gene program initialization improves average PCC by 69.6% and 58% on the two tasks, respectively, compared to using the foundation models alone. This indicates that EP-Booster is more effective at reconstructing a task-relevant latent space, providing a more suitable feature representation for gene expression prediction. As PCA focuses on directions of maximal variance, it potentially discards biologically informative but low-variance features. However, EP-Booster leverages biologically informed prototypes to guide latent space reconstruction. This approach improves the preservation of relevant information about gene expression while simultaneously filtering out noise and redundancy.

### Biologically Relevang Gene Expression Signatures Can Be Predicted

Gene expression signatures consist of sets of genes whose coordinated expression patterns characterize specific biological states or disease phenotypes. They are widely used in disease classification, prognosis, and treatment response prediction, making their accurate inference from histopathology images an important and clinically relevant task. Accordingly, we evaluate our method on two representative gene expression signature sets: breast cancer signatures [15, 16] and melanocyte identity signatures [17, 18]. In the IDC dataset, 16 breast cancer signatures are predicted using Virchow 2 alone as well as our EP-Booster with both random and gene program–guided initialization. As shown in Figure 3a and c, EP-Booster consistently outperforms Virchow 2, demonstrating its effectiveness in predicting clinically relevant gene expression signatures. We further evaluate melanocyte identity signature prediction on the SKCM dataset. As shown in Figure 3b and d, EP-Booster achieves substantially better performance than Virchow 2. Notably, using gene program–guided initialization consistently outperforms random initialization in this setting, suggesting that incorporating biological prior knowledge is particularly beneficial when predicting structured gene expression signatures.

**Fig. 3.**
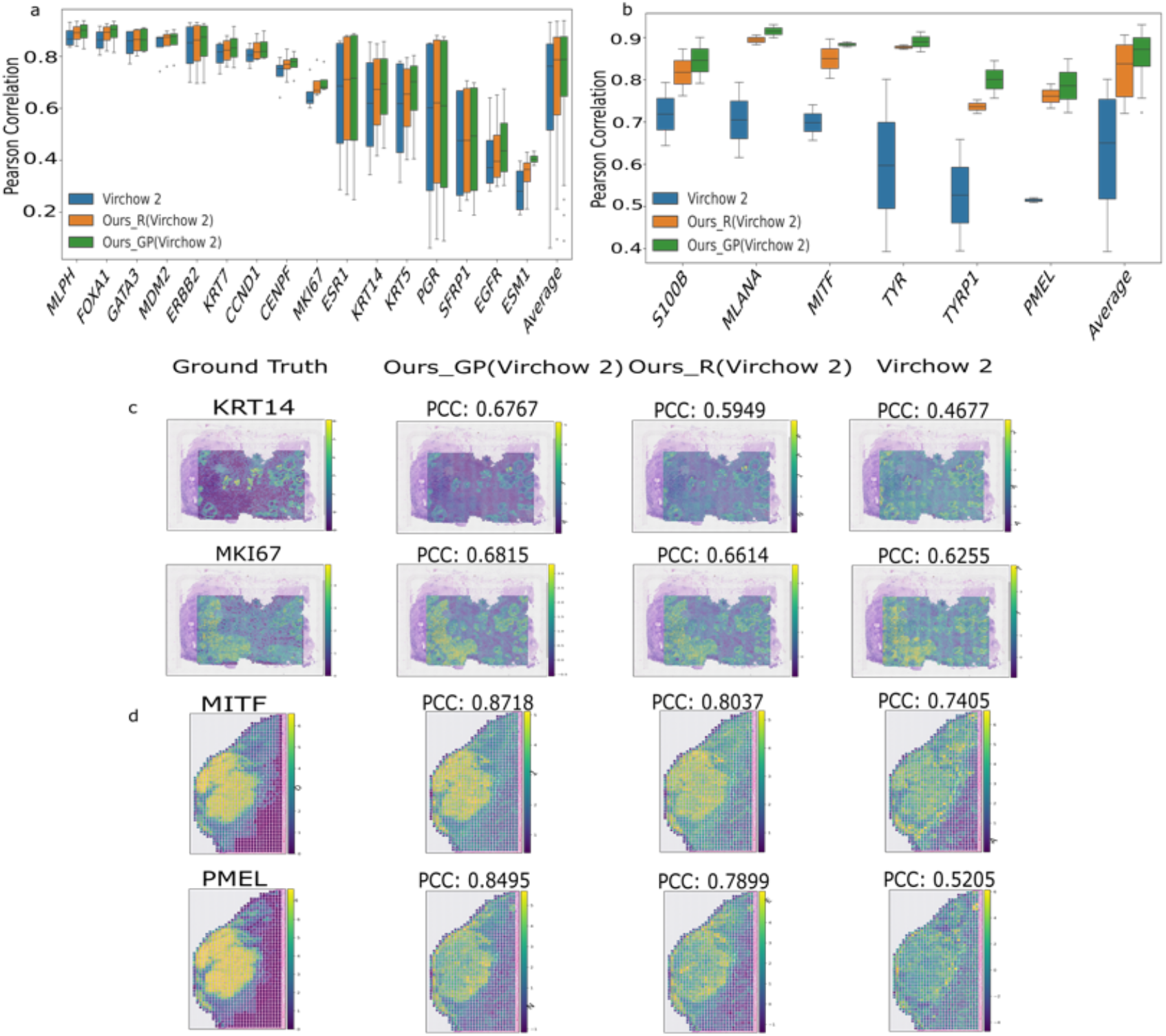
Superior performance of EP-Booster compared with foundation models. **a**, Breast cancer signature prediction results of EP-Booster and Virchow 2 on the IDC dataset, showing four representative samples and 16 genes. **b**, Melanocyte identity signature prediction results on the SKCM dataset, showing two representative samples and 6 genes. **c**, Predicted spatial distributions of representative breast cancer signatures (KRT14, MKI67) in the IDC dataset. **d**, Predicted spatial distributions of representative melanocyte identity signatures (MITF, PMEL) in the SKCM dataset.

### Evaluation on Clinically Relevant Tasks

To externally validate the clinical potential of EP-Booster, we conducted survival analysis and drug response prediction on the TCGA and transNEO datasets. For survival analysis, EP-Booster–predicted gene expression showed reasonable consistency with bulk RNA-seq data, as reflected by gene-wise PCC (Figure 4a). Using the top five genes selected via the univariate Cox regression model, we performed survival analysis on TCGA samples with matched clinical data. Kaplan–Meier analysis demonstrated significant stratification between high- and low-risk groups using both predicted and actual gene expression (Figures 4b, c). Notably, predicted expression achieved slightly improved risk stratification performance relative to bulk RNA-seq, suggesting that EP-Booster captures prognostically relevant signals from histopathological morphology.

**Fig. 4.**
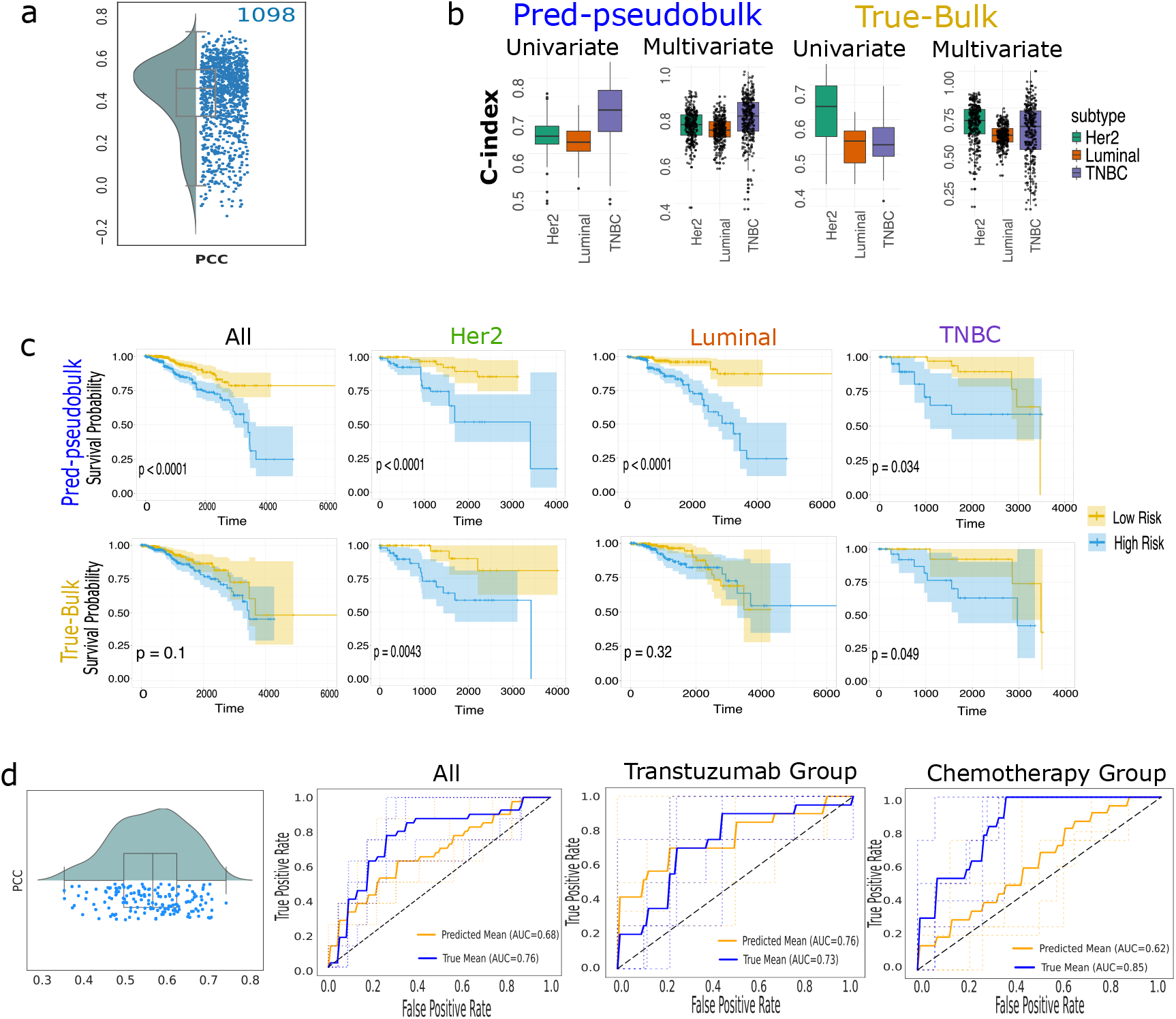
EP-Booster Predictions for H&E images enable stratification of patient survival and drug response. **a**, Box plots showing gene-wise PCC per sample, computed between EP-Booster–predicted gene expression from TCGA H&E images and matched bulk RNA-seq data. **b**, Performance of prognostic models evaluated using the concordance index (C-index) in both univariate and multivariate Cox regression models, based on predicted (pred-pseudobulk) and actual (true-bulk) gene expression. **c**, Kaplan–Meier survival analysis stratifying patients into high- and low-risk groups using either bulk RNA-seq (true-bulk) or EP-Booster-predicted gene expression (pred-pseudobulk). **d**, Box plots showing gene-wise PCC per sample between predicted and bulk gene expression, together with AUC comparisons for drug response prediction using four machine learning models under different data settings.

For drug response prediction, the ensemble model (more details in Section Implementation Details) based on predicted gene expression achieved consistent performance on the transNEO cohort (Figure 4d), with an overall AUC of 0.68 compared to 0.76 using actual gene expression. Subgroup analysis further confirmed its robustness, with AUCs of 0.76 for trastuzumab (vs. 0.73 using actual expression) and 0.62 for chemotherapy, where consistent discriminative trends between responders and non-responders were still observed.

### Contribution of EP-Booster components and gene program initialization

#### Ablation Analysis of EP-Booster Components

In the ablation studies, we evaluate the contribution of each component within EP-Booster. The results are summarized in Table 2. We employ a Ridge regression model as the baseline for comparison. Each component is incrementally added to assess its impact on the overall performance, allowing us to isolate the effectiveness of the prototype-guided cross-attention module, gating mechanisms, gene program initialization, and two loss functions. We began by incorporating a prototype-guided cross-attention module into the baseline model, which led to significant performance improvements (PCC gain of 110.6%). Next, we independently added the *g*_*prototype*_ and *g*_*dim*_. The results show that adding *g*_*dim*_ yields greater performance gains than *g*_*prototype*_. When both gating mechanisms were integrated into the prototype-guided cross-attention module, the resulting model is called EP-Booster with random initialization. It further improves performance, achieving PCC gain of 1.6%. Then, initializing the prototypes using gene programs (GPs) led to further performance gains, demonstrating the effectiveness of incorporating biological priors into prototype construction. Finally, we evaluate the contributions of each loss function. Using the *L*_*Recon*_ and *L*_*Reg*_ individually produces slightly lower performance than their combined use. This likely reflects their complementary objectives: *L*_*Recon*_ ensures the EP-Booster embedding retains information from the original image embedding, while *L*_*Reg*_ enforces gene expression relevance in the embedding.

**Table 2.**
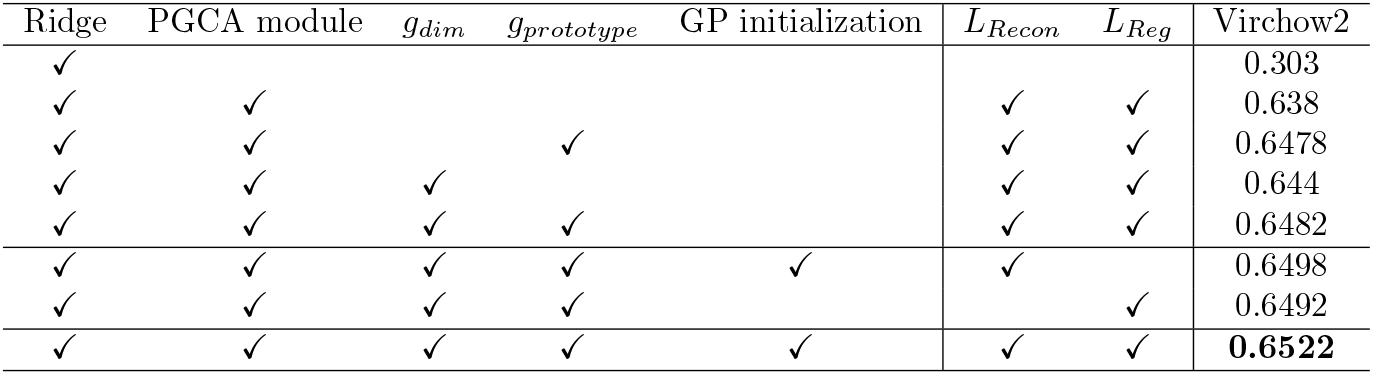
Ablation analysis of the proposed method on the SKCM dataset from the HEST benchmark[10]. Reported results are the mean values of PCC obtained from cross-validation. PGCA module: the prototype-guided cross-attention module.

#### Analysis of Embeddings across Models Shows Superior Formation of the Latent Space

We conduct a comparative analysis of the original Virchow2 embeddings, PCA-reduced embeddings, and EP-Booster–enhanced embeddings using t-SNE visualization and k-means clustering (k = 10), as shown in Figure 5a. The original embeddings appear highly dispersed in the latent space with substantial cluster overlap, resulting in fragmented spatial patterns that obscure clear anatomical structures. Applying PCA partially compresses the feature space and improves cluster compactness, but cluster boundaries remain indistinct and spatial coherence is still limited. In contrast, EP-Booster–enhanced embeddings exhibit substantially improved structure: clusters show higher internal density and clearer separation in the t-SNE space. This improved representation also leads to stronger spatial consistency, where points sharing cluster labels display clear spatial aggregation aligned with the underlying tissue architecture. By reducing redundant noise and enhancing feature separability, EP-Booster produces more structured embeddings that support more accurate downstream gene expression prediction.

**Fig. 5.**
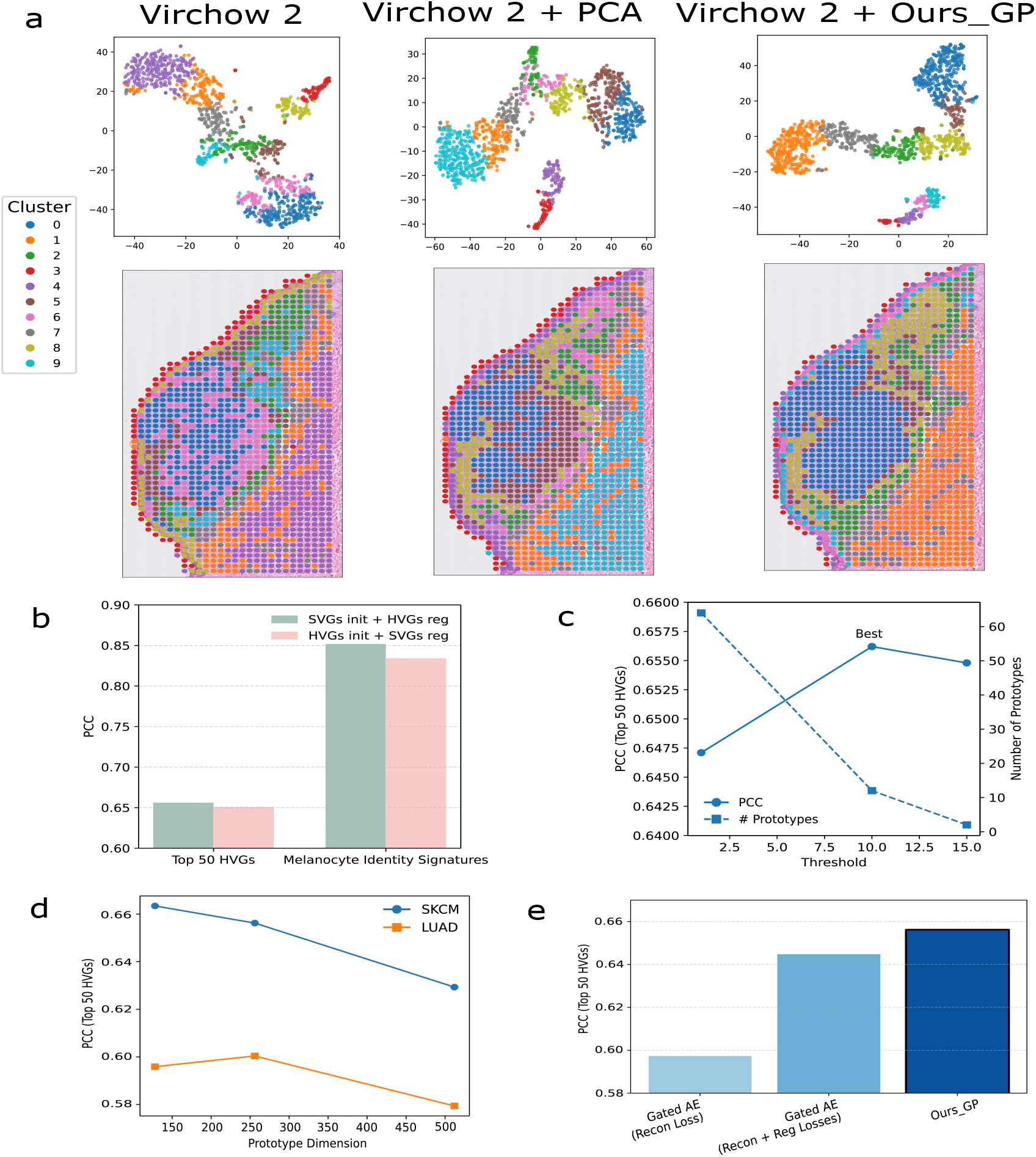
Ablation studies of EP-Booster components and gene program initialization. **a**, T-SNE and spatial visualizations with k-means clustering of different embedding representations on the SKCM dataset. **b**, Comparison of two gene list utilization strategies evaluated across two gene expression prediction tasks on the SKCM dataset. **c**, Sensitivity analysis of the model performance and the number of prototypes by varying the threshold used in gene program–based initialization on the SKCM dataset. **d**, Performance comparison across different prototype dimensionalities on the SKCM and LUAD datasets for the top 50 HVG prediction task. **e**, Comparison between a simple MLP-based autoencoder (with and without biological supervision) and EP-Booster on the top 50 HVG prediction task on the SKCM dataset. For all ablation studies, Virchow 2 is used as the foundation model.

#### The Prototype Approach Effectivly Integrates Gene Programs and Biological Information

We further conduct ablation studies to examine the effects of prototype configuration (including number and dimensionality), different gene sets used for prototype initialization and supervised regression, and the contribution of biological information. The results are summarized in Figure 5(b–e).

##### Gene set utilization strategy

The motivation for employing two distinct gene sets, such as SVGs and HVGs, arises from the different functional roles of prototype initialization and supervised regression. SVGs are used to construct gene programs for prototype initialization because they exhibit spatially coherent expression patterns and provide biologically meaningful priors that align well with the morphological structures captured in H&E image embeddings. In contrast, not all HVGs encode spatial information. HVGs are instead employed for supervised regression, as their high expression variability typically yields clearer gradients and stronger supervision signals. To validate this design choice, we perform additional experiments on the SKCM dataset by swapping the roles of SVGs and HVGs for both the top 50 HVGs and melanocyte identity signatures prediction tasks (Figure 5b). The results show that using SVGs for prototype initialization and HVGs for regression slightly outperforms the flipped configuration, supporting our design choice.

##### Sensitivity to gene program initialization threshold

We next investigate how model performance and the number of prototypes are affected by varying the threshold used in gene program initialization (Figure 5c). When the threshold is set to 1, meaning a gene program is activated if it shares at least one SVG with the selected genes, a total of 64 gene programs are included. This setting leads to decreased performance compared to a threshold of 10, likely because gene programs containing only a single SVG introduce noise rather than providing informative priors. Conversely, increasing the threshold to 15 retains only two gene programs, resulting in overly restrictive prior information and insufficient biological guidance, which again slightly degrades performance. These results indicate that the threshold plays a critical role in balancing the richness and noise of biological priors used for prototype initialization.

##### Prototype dimensionality

We further perform ablation studies on prototype dimensionality for the top 50 HVGs prediction task (Figure 5d). The results reveal that different cancer types exhibit distinct performance trends under the Virchow 2 foundation model as the prototype dimensionality varies. Accordingly, the default prototype dimension is selected by jointly considering all datasets for each foundation model, aiming for a balanced and robust configuration.

##### Effect of biological information injection

Finally, to isolate the contribution of biological information, including both biological priors and biological supervision, we replace EP-Booster with a simple MLP-based gated autoencoder. This baseline shares the same decoder and regression head as EP-Booster, and its encoder is parameter-matched to the prototype-guided cross-attention module. We evaluate this variant on the SKCM dataset for the top 50 HVGs prediction task. As shown in Figure 5e, EP-Booster consistently outperforms the gated autoencoder, demonstrating that the observed performance gains arise from the incorporation of biological information rather than from increased model capacity.

#### Analysis of Gene Program and Interpretability

For different disease types, the gene programs differ. We extract gene programs list for SKCM and LUAD datasets, which is shown in Table S2. The Table shows that different cancer types have different numbers and types of pathways selected for prototype initialization. Lung cancer is dominated by classical oncogenic signalling, including EGFR, ALK, KRAS–RAF–MEK, and AKT–mTOR, leading to enrichment of genes in RTK activation and cell-cycle progression [19, 20]. In contrast, skin cancer, especially melanoma, shows more of stemness and developmental pathways such as Wnt/LEF1 and SHH, together with RAF activation, reflecting more genes in dedifferentiated and MAPK-driven profiles [21].

In addition, to validate the biological basis of the model predictions, we performed pathway-level interpretability analyses. Figure 6 shows the pathway contribution decomposition for a representative image patch (Patch 1415), where TYRP1 (Figure 6a) shows a high ground-truth expression (5.68) and is accurately predicted by the model (5.41). This prediction is primarily driven by the cAMP signalling pathway (CAMP UP.V1 DN), followed by the Wnt//*β* -catenin pathway (LEF1 UP.V1 UP), both of which are established upstream regulators of MITF, the master transcription factor governing melanogenesis in melanoma. Extending this analysis to additional melanogenesis-related markers within the same patch (Figure 6 (b-d)), the model achieved close agreement between predicted and true expression values for MITF (4.42 vs. 4.32), PMEL (5.08 vs. 6.15) and S100B (4.27 vs. 5.52). Notably, pathway contribution decomposition across all four genes revealed a highly consistent hierarchical pattern, with the cAMP and Wnt/*β* -catenin pathways consistently ranked as the top two contributors. This attribution precisely mirrors the canonical melanocyte regulatory axis, in which cAMP and Wnt signalling cooperatively activate MITF, which in turn drives the expression of TYRP1 and PMEL. These results demonstrate that the model reconstructs biologically meaningful representations directly from histopathological morphology, providing strong evidence that its interpretability is grounded in underlying biological mechanisms. Results of the interpretability analysis on the IDC dataset are shown in Figure S3.

**Fig. 6.**
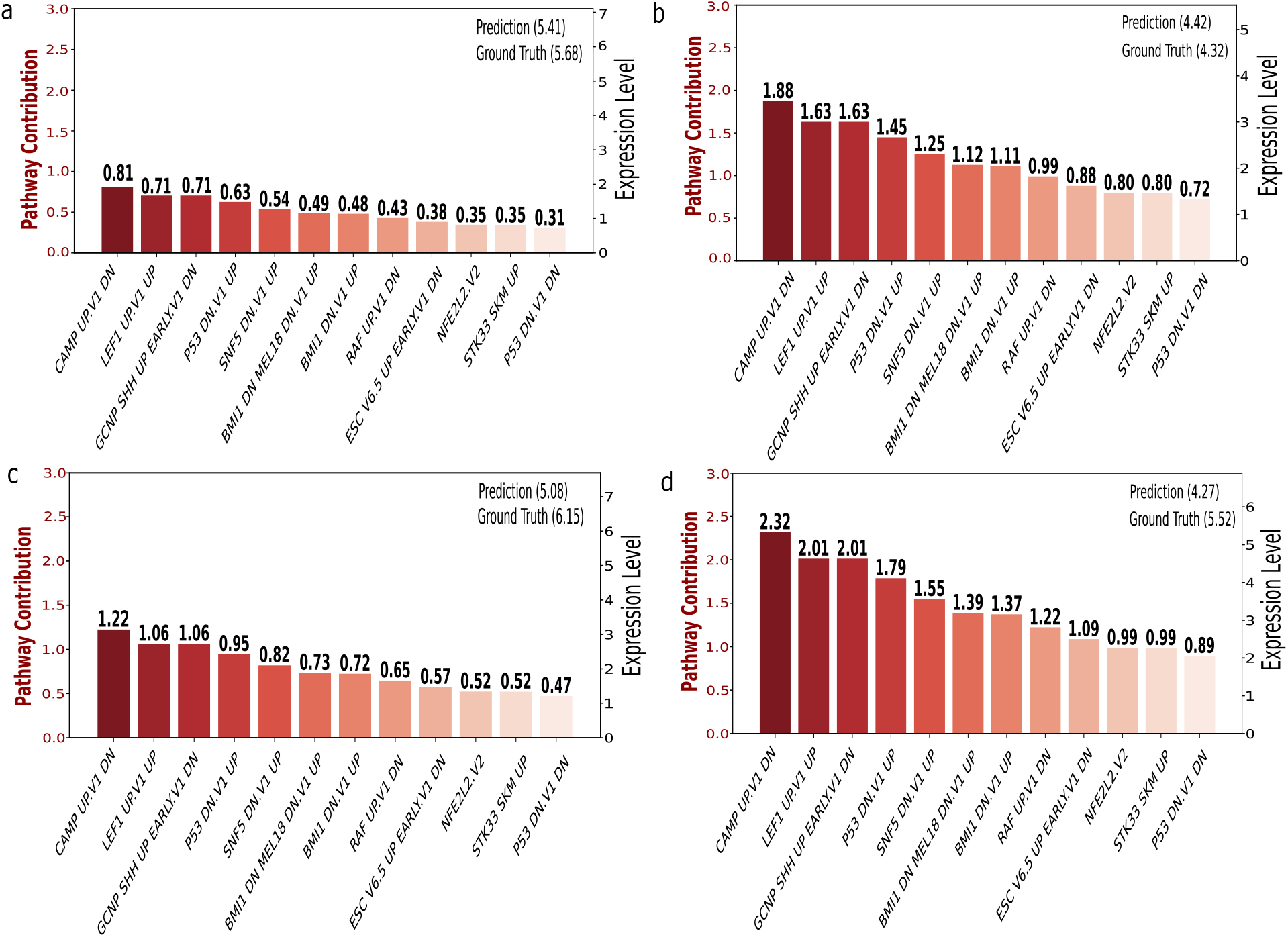
Interpretability analysis of marker gene predictions in the SKCM dataset. Contribution analysis for representative marker genes: **a** TYRP1, **b** MITF, **c** PMEL and **d** S100B. These genes are key markers of melanocyte differentiation and melanoma progression.

## Discussion

In this study, we propose EP-Booster, a lightweight refinement module that reconstructs latent representations from existing pathology foundation models through biologically guided prototype learning to improve gene expression prediction. Importantly, we leverage pathway-level interpretability to associate model predictions with biologically relevant pathways derived from gene program–based initialization, thereby improving interpretability. EP-Booster also demonstrates robust performance across different ST platforms and tissue types while enhancing the task-specific representational capacity of foundation models, highlighting its applicability to gene expression signatures, survival analysis, and drug response prediction. We expect that EP-Booster, which reconstructs foundation model image embeddings by incorporating biological knowledge, can offer new insights into leveraging foundation models and H&E images for computational pathology.

Our EP-Booster is conceptually different from domain adaptation, multi-modal alignment, and feature recalibration. First, unlike domain adaptation, which seeks to mitigate distribution shifts and learn domain-invariant features via aligning distribution approaches, our method focuses on semantically adapting visual features to molecular profiles, addressing modality-level semantic differences rather than distribution mismatch. Second, compared with multi-modal alignment methods such as contrastive learning [22], which typically embed two modalities into a shared latent space for bidirectional retrieval or generation, our method performs a unidirectional refinement of image embeddings, where SVGs and HVGs serve as supervised signals to guide EP-Booster in refining the embeddings rather than mapping image and gene expression features into a shared representation space. This makes the embeddings more suitable for gene expression prediction. Finally, unlike classic feature recalibration approaches such as SENet [23], which compute channel weights using internal feature statistics, our method reconstructs embeddings by modelling the correlation between image features and biologically informed prototypes, providing improved interpretability. In addition, our attention and gating design differ from recent gated attention mechanisms [24] that apply head-specific sigmoid gates to modulate scaled dot-product attention outputs. While such approaches primarily introduce non-linearity and stabilize attention dynamics at the token level, EP-Booster employs prototype-guided cross-attention with gating over both prototypes and latent dimensions, enabling biologically structured refinement of image embeddings. A systematic comparison with these gating strategies will be explored in future work.

Despite the advantages of EP-Booster in gene expression prediction, several aspects remain for further improvement. First, the current gene program construction relies on SVGs shared across all training samples, which may exclude tissue-specific SVGs and weaken gene program-guided prototype initialization. Future work could explore alternative SVG selection strategies, such as sample-specific SVG identification combined with sample-wise training. Second, each prototype is currently statically associated with a biological pathway. However, the morphological expression of the same biological process can vary substantially across tissue regions and microenvironments. Introducing conditional prototypes, where sample-specific image features modulate prototype initialization, could enable adaptive focus on diverse morphological patterns of the same pathway, thereby improving generalization without retraining.

## Methods

### Prototype-guided Cross-attention Module

The core of EP-Booster is a lightweight prototype-guided cross-attention module composed of: a projection layer, a learnable prototype set *P* and a multi-head cross-attention layer. Since the dimension of *Z*_img_ can vary across foundation models, we first apply a projection layer to align it with the prototype dimension *L*_2_. The multi-head cross-attention module then computes prototype-aware representations 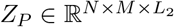 by taking **Proj**(*Z*_img_) and *P* as inputs, as follows:

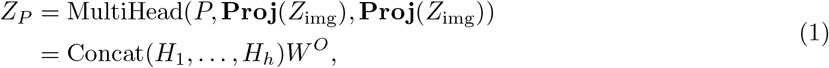

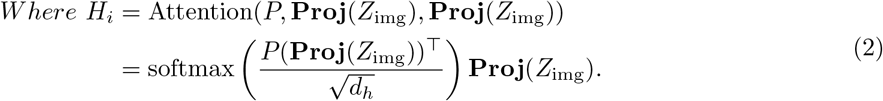

Here, 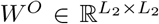 is the output projection matrix, *d*_*h*_ = *L*_2_*/h* is the head dimension, and **Proj**(·) is the projection layer. This design allows each prototype to attend to distinct semantic subspaces of the image embeddings, improving cross-modal alignment.

### Gating Mechanisms

To improve representation quality and interpretability, EP-Booster incorporates two learnable gating mechanisms:

#### Prototype Gate

Each prototype is associated with a learnable scalar gate *g*_prototype_ ∈ [0, 1]^*M*^. The gated prototype output is:

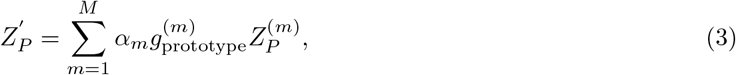

where *α* denotes the attention scores from the prototype-based cross-attention layer and *M* is the number of prototypes.

#### Latent Dimension Gate

A second gate 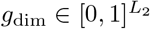 is applied to control each latent dimension:

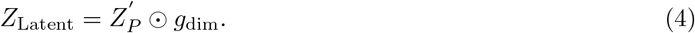

where ⊙ represents element-wise multiplication. These mechanisms selectively suppress uninformative prototypes and dimensions, enhancing both model compactness and task-specificity.

### Decoder and Regression Head

To ensure that *Z*_Latent_ preserves essential visual information from *Z*_img_ while being predictive of gene expression, we introduce a decoder that reconstructs *Z*_img_ from *Z*_Latent_, supervised with a reconstruction loss *L*_Recon_. A regression head that maps *Z*_Latent_ to the HVGs selected from the training set, with prediction supervised by a regression loss *L*_Reg_. Together, these losses enforce that *Z*_Latent_ is both faithful to visual input and biologically meaningful, driving improved generalization on downstream gene expression prediction tasks.

### Prototype Initialization Strategies to Improve Performance

In EP-Booster, prototypes play a central role in transforming *Z*_img_ into the refined latent space *Z*_latent_. The strategy for initializing these prototypes is therefore an important design choice. To this end, we propose a prototype initialization strategy that supports two modes: random initialization and gene program-guided initialization. This strategy enables flexibility in leveraging purely data-driven or biologically informed priors during model training.

### Random Initialization

The baseline initialization strategy involves sampling the learnable prototypes from a Gaussian distribution, given by:

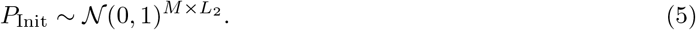

To ensure that the resulting latent representation *Z*_Latent_, constructed from these randomly initialized prototypes and the image embeddings *Z*_img_, encodes not only essential visual features but also biologically meaningful information related to gene expression, we employ both *G*^HVG^ and *Z*_img_ as supervisory signals. These guide the prototypes to capture visual-biological correspondences during training. This strategy enables the model to learn biologically relevant abstractions in a data-driven manner, without requiring prior domain knowledge for prototype initialization. However, random initialization lacks semantic grounding and makes it difficult to interpret what each prototype represents, thereby limiting the interpretability of the learned latent space. To address this limitation, we propose an alternative gene program–guided initialization strategy, driven by spatially variable genes (SVGs).

### Gene Program-Guided Initialization

A gene program is a set of co-expressed genes associated with a specific biological process. It is typically constructed by integrating biological pathways with data-driven co-expression patterns to identify biologically meaningful and context-specific gene modules [25]. Existing gene program construction methods, such as ExpiMap [26], primarily rely on filtering highly variable genes (HVGs). While HVGs capture genes with significant expression variation across samples, they often overlook the spatial organization of tissues. In contrast, spatially variable genes (SVGs) capture expression differences across distinct tissue locations, thus better reflecting spatial heterogeneity.

Inspired by this, we propose a prototype initialization strategy driven by spatially variable genes (SVGs) and enriched with biological priors at the pathway level. This strategy enables a semantically meaningful initialization for prototypes, improving interpretability and providing a strong starting point for subsequent training. Specifically, we first collect a set of biological pathways *GP* = { *GP*_1_, *GP*_2_,…} from a pathway database such as MSigDB [27]. These pathways are filtered by intersecting them with the SVG set *G*^SVG^, retaining only those that satisfy the condition that the number of genes | *G*^SVG^ ∩ *GP* | ≥ 10. Based on the filtered results, we build the pathway-SVG binary matrix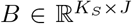, where *K*_*S*_ is the number of SVGs and *J* is the number of selected pathways. Then, the gene program matrix is defined as:

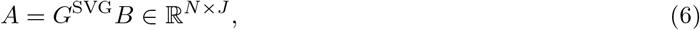

*N* denotes the total number of image patches. Then, we apply Sigmoid activation to each gene program 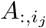, followed by normalization to compute the spatial weights:

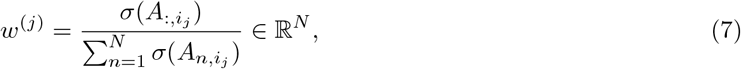

where *σ* (·) denotes the Sigmoid activation function. Using these weights, we define the *j*-th prototype as the weighted sum of SVG expression across spatial positions:

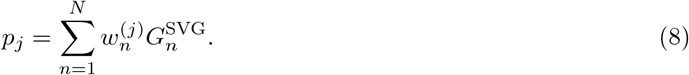

Finally, by stacking all prototypes and passing them through a projection layer, we obtain the initialized prototype matrix:

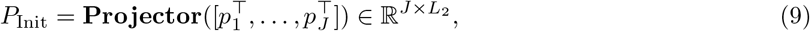

A key advantage of this initialization strategy is that, unlike random initialization, it eliminates the need to manually specify the number of prototypes by directly leveraging gene programs as semantically meaningful prototypes (*i*.*e*.,*M* = *J*). Moreover, since each prototype corresponds to a filtered pathway, the model gains clear biological interpretability by linking prototypes to known pathways.

### Joint Reconstruction-Regression Optimization

In our approach, EP-Booster is optimized using a combination of a reconstruction loss and a regression loss, defined as follows:

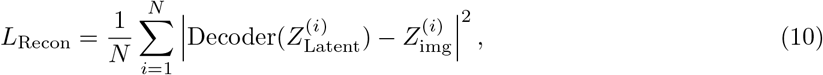

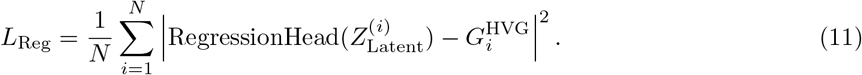

The overall training objective is the sum of these two components:

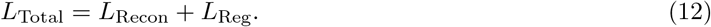

This joint optimization strategy encourages the model to learn latent representations that are not only faithful to the original image embeddings but also predictive of biologically meaningful gene expression patterns.

### Pathway-Level Interpretability

To enhance the interpretability of EP-Booster, we propose a pathway-level interpretability method based on gene program–guided initialization and the prototype-guided cross-attention module. This design enables model predictions to be explicitly traced back to biologically meaningful pathways. As described in Sections Gene Program-Guided Initialization and Prototype-guided Cross-attention Module, pathway information is encoded into the prototypes through gene program–guided initialization. During inference, the prototype-guided cross-attention module aligns the image embeddings with the prototypes, producing attention scores that reflect the relevance of each pathway, while the prototype gate *g*_prototype_ assigns a weight to each pathway. As a result, the reconstructed latent embedding incorporates pathway information and can be expressed as:

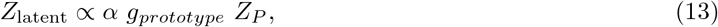

where *α* denotes the attention scores. The latent representation *Z*_latent_ is subsequently used as input to a ridge regression model for gene expression prediction:

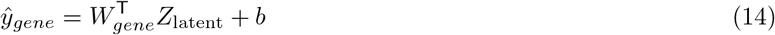

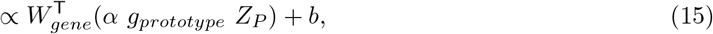

where *W*_*gene*_ denotes the regression weights associated with gene *g*. Under this formulation, the contribution of each prototype, corresponding to a biological pathway, to the prediction of a gene can be quantified as:

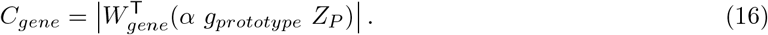

In this work, we focus on assessing the magnitude of pathway contributions to the predicted gene expression. Therefore, absolute values are used to reflect the overall influence of each pathway, irrespective of direction.

### Diverse Datasets for Model Training and Evaluation

We evaluated our method across multiple spatial transcriptomics (ST) and clinical cohorts spanning diverse cancer types, ST platforms, and prediction tasks. For the ST cohort, we used four cancer types from the HEST benchmark [10], including IDC (invasive ductal carcinoma), SKCM (skin cutaneous melanoma), PRAD (prostate adenocarcinoma), and LUAD (lung adenocarcinoma). This benchmark provides standardized, high-quality datasets generated using different spatial transcriptomics technologies, enabling evaluation under cross-platform variability. The Xenium-based cohorts (IDC, SKCM, and LUAD) comprise a total of 8 tissue sections (n = 4, 2, and 2, respectively), where each tissue section corresponds to a distinct patient (i.e., 8 patients in total). These datasets capture heterogeneous tumor microenvironments across breast, skin, and lung cancers. We followed [10] and adopted cross-validation to assess model generalization. The PRAD cohort, generated using Visium technology, includes 23 tissue sections derived from 2 patients, providing a dataset with repeated spatial measurements per patient. Evaluation was conducted using a leave-one-out strategy [9], enabling assessment of generalization under limited patient diversity.

To further evaluate clinical utility, we assessed survival analysis on The Cancer Genome Atlas (TCGA) cohort, comprising n = 670 patients with matched histopathology images (n = 1,034), bulk RNA-seq profiles, and survival outcomes. Whole-slide H&E images were obtained from TCIA [28], and corresponding molecular and clinical data were collected from the TCGA portal^1^. Following data quality control procedures (detailed in Section Data Preprocessing), a subset of high-quality breast cancer samples was retained for analysis. In addition, we evaluated drug response prediction on the transNEO dataset [29], which comprises n = 160 breast cancer patients with matched histopathology slides, RNA sequencing (RNA-seq) data, and treatment response information. Patients are stratified into responder and non-responder groups based on pathological complete response (pCR), enabling clinically meaningful evaluation of treatment outcomes. In our analysis, we further considered treatment-specific subgroups, including trastuzumab-treated and chemotherapy-treated patients, to assess model performance across different therapeutic settings.

### Data Preprocessing

We follow the HEST benchmark [10] for data preprocessing across four cancer types in the ST datasets. Highly variable genes (HVGs) and spatially variable genes (SVGs) are selected only from training data. For datasets with multiple training samples, we retain genes that are commonly identified as HVGs or SVGs across all samples. For the TCGA dataset, we obtained 3,066 H&E images from TCIA. 1,532 images were excluded due to the absence of matched genomic expression profiles and clinical data. The remaining 1,534 images were subjected to quality control using the approach described in [30], resulting in the removal of 500 low-quality images. Ultimately, 1,034 breast cancer images from 670 patients were retained for survival analysis.

### Gene List Selection Strategy for Training

In our method, two biologically distinct gene sets are utilized to initialize and train the prototypes: SVGs and HVGs. SVGs are genes whose expression levels exhibit significant variation across spatial locations within a tissue, capturing spatial heterogeneity. We use Squidpy [31] to identify the top 200 SVGs from the training set, which are employed for prototype initialization. HVGs represent genes with high expression variability across samples and are commonly used to capture key data structures. We select the top 200 HVGs from the training data using Scanpy [32] and use them for prototype training. We found no significant performance change when varying the number of SVGs or HVGs.

### Foundation Models

In this study, we evaluate our method using nine foundation models across different experiments: ResNet-50 [33], CTransPath [34], Phikon [35], CONCH [8], GigaPath [36], UNI [7], Virchow [37], Virchow 2 [38], and H-Optimus-0 [39].

### Implementation Details

Our model is implemented using the PyTorch framework and executed on an NVIDIA L40 GPU (48GB memory). We use Adam optimizer to train our model for 50 epochs. The batch size is 128. The hyperparameters related to the prototype (the dimension and number) and the learning rate are adjusted based on the choice of foundation model and validation dataset. The learning rate was selected from the range [0.00001, 0.005]. The prototype dimension was chosen from {128, 256, 512}, and the number of prototypes from {16, 32, 64}. Detailed hyperparameter configurations for each experiment are provided in the Appendix 1. For evaluation, we follow the HEST benchmark and first use the **top 50 HVGs** to assess performance on the IDC, SKCM, and LUAD datasets. Second, we adopt the **high mean and high variability gene (HMHVG)** list, as defined in Stem [9], to evaluate all methods on the PRAD dataset. Additionally, we evaluate gene expression signatures, including **breast cancer signatures** on the IDC dataset and **melanocyte identity signatures** on the SKCM dataset, demonstrating the effectiveness of EP-Booster in predicting clinically important biomarkers. To measure gene expression prediction performance, we use the Pearson Correlation Coefficient (PCC) as an evaluation metric.

In addition, we evaluate the clinical applicability of EP-Booster through external tasks, including survival analysis and drug response prediction. Our model was trained on the IDC dataset from the HEST benchmark using the top 200 HVGs identified from the IDC cohort, and applied to predict gene expression in the TCGA dataset for survival analysis and in the transNEO dataset for drug response prediction. For survival analysis, predicted gene expression was compared with TCGA bulk gene expression data. Univariate Cox regression models were constructed for each gene and evaluated using the concordance index (C-index), followed by multivariate models built from the top five genes and assessed via repeated three-fold cross-validation. Patients were split into low- and high-risk groups, with analyses performed separately for each molecular subtype. For drug response prediction, four classical machine learning models (logistic regression, support vector machines, k-nearest neighbours, and random forests) were trained on the predicted gene expression via a nested 5-fold cross-validation approach. Within each training fold, gene expression was normalised, the top 100 genes were selected using ANOVA F-values, and final predictions were obtained by averaging model outputs. Model performance was evaluated using AUC.

## Data Availability

The data underlying this study are available from multiple sources. The HEST-1k dataset is publicly accessible at https://huggingface.co/datasets/MahmoodLab/hest. For the TCGA dataset, H&E images were obtained from TCIA [28], and matched bulk gene expression and clinical data were collected from the TCGA data portal (https://www.cancer.gov/ccg/research/genome-sequencing/tcga). The transNEO dataset is available in [29].

## Code Availability

All code is available at https://github.com/GenomicsMachineLearning/NCSTAR/tree/main/EP-Booster.

## Author contributions statement

Q.N. and C.L. conceived and designed the study. C.L. and Q.N. jointly designed the model. C.L. implemented the model and performed all experiments. Q.N. supervised the project and designed experiments related to clinical application. C.L. and Q.N. analyzed and interpreted the data. All authors contributed to the writing of the manuscript.

## Acknowledgments

The authors thank the anonymous reviewers for their valuable suggestions.

## Supplemental Materials

## Supplementary Figures

**Fig. S1.**
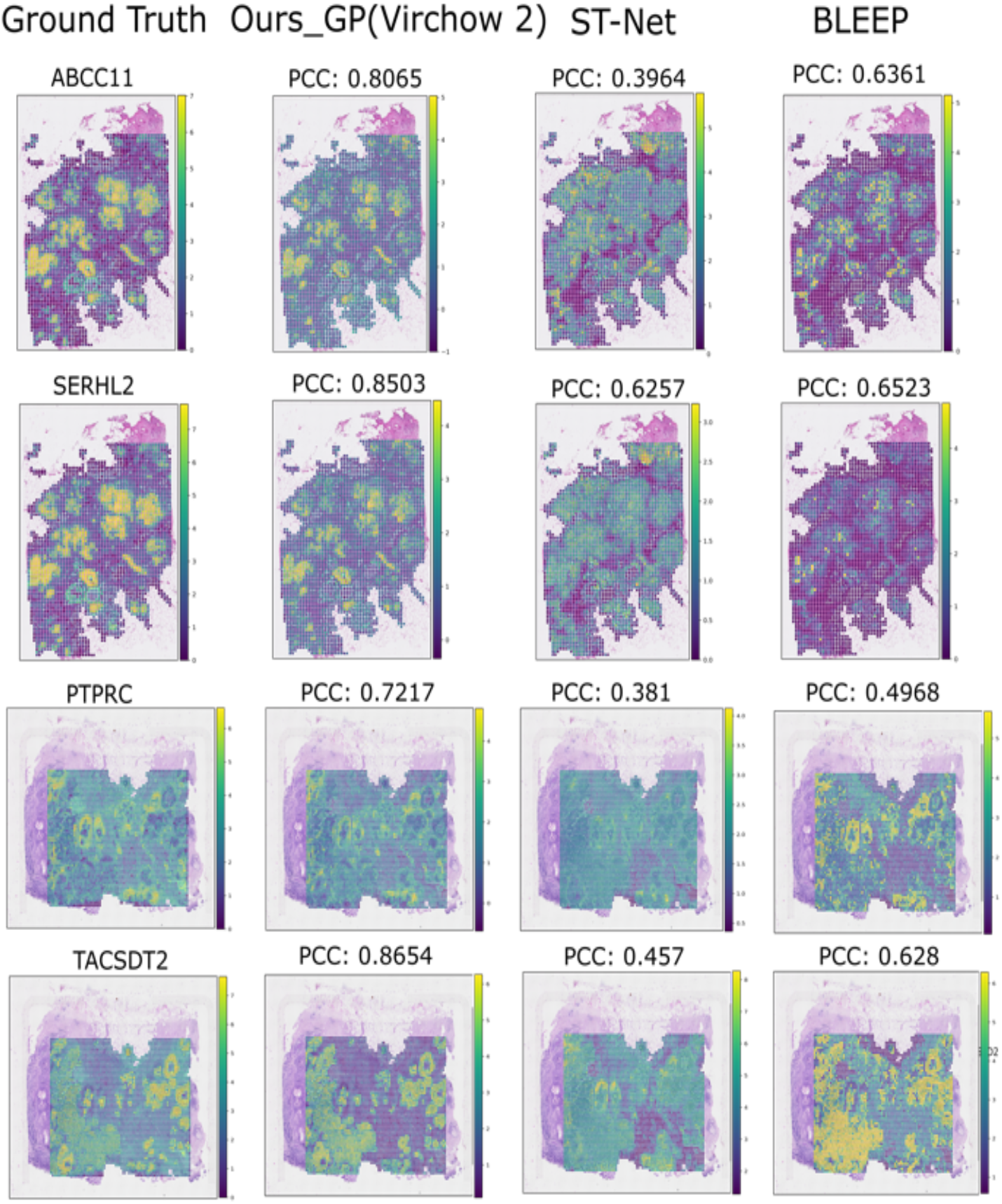
Visual comparison of spatial expression predictions for highly variable genes in the IDC dataset. Spatial expression patterns of four representative highly variable genes (ABCC11, SERHL2, PTPRC and TACSDT2) are shown for ground truth and predictions from different models (Ours_GP, ST-Net and BLEEP). Pearson correlation coefficients (PCCs) are indicated above each panel. Gene expression values are log-normalized, and the colour scale represents expression intensity (yellow, high; purple, low).

**Fig. S2.**
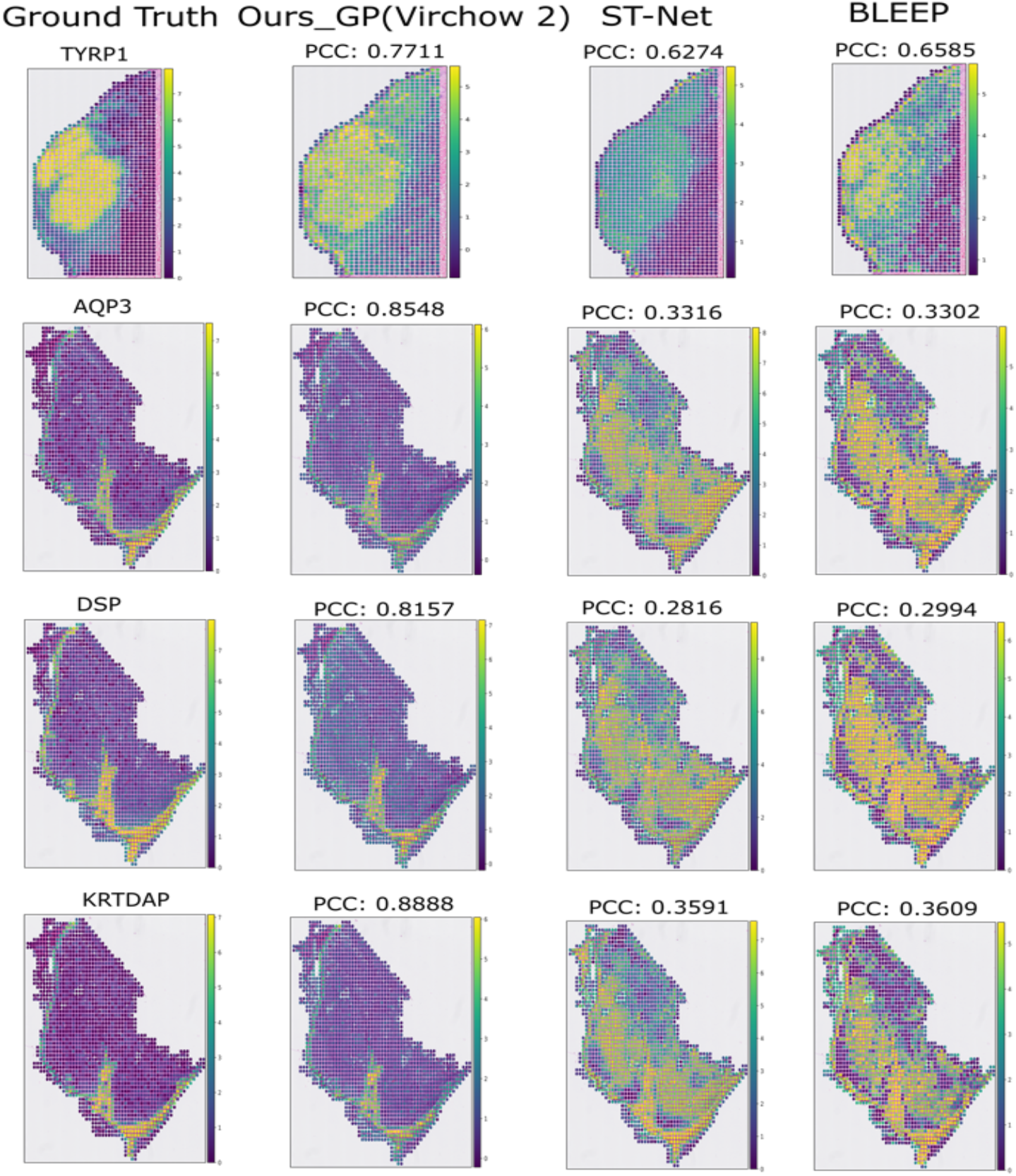
Visual comparison of spatial expression predictions for highly variable genes in the SKCM dataset. Spatial expression patterns of four representative highly variable genes (TYRP1, AQP3, DSP and KRTDAP) are shown for ground truth and predictions from different models (Ours_GP, ST-Net and BLEEP). Pearson correlation coefficients (PCCs) are indicated above each panel. Gene expression values are log-normalized, and the colour scale represents expression intensity (yellow, high; purple, low).

**Fig. S3.**
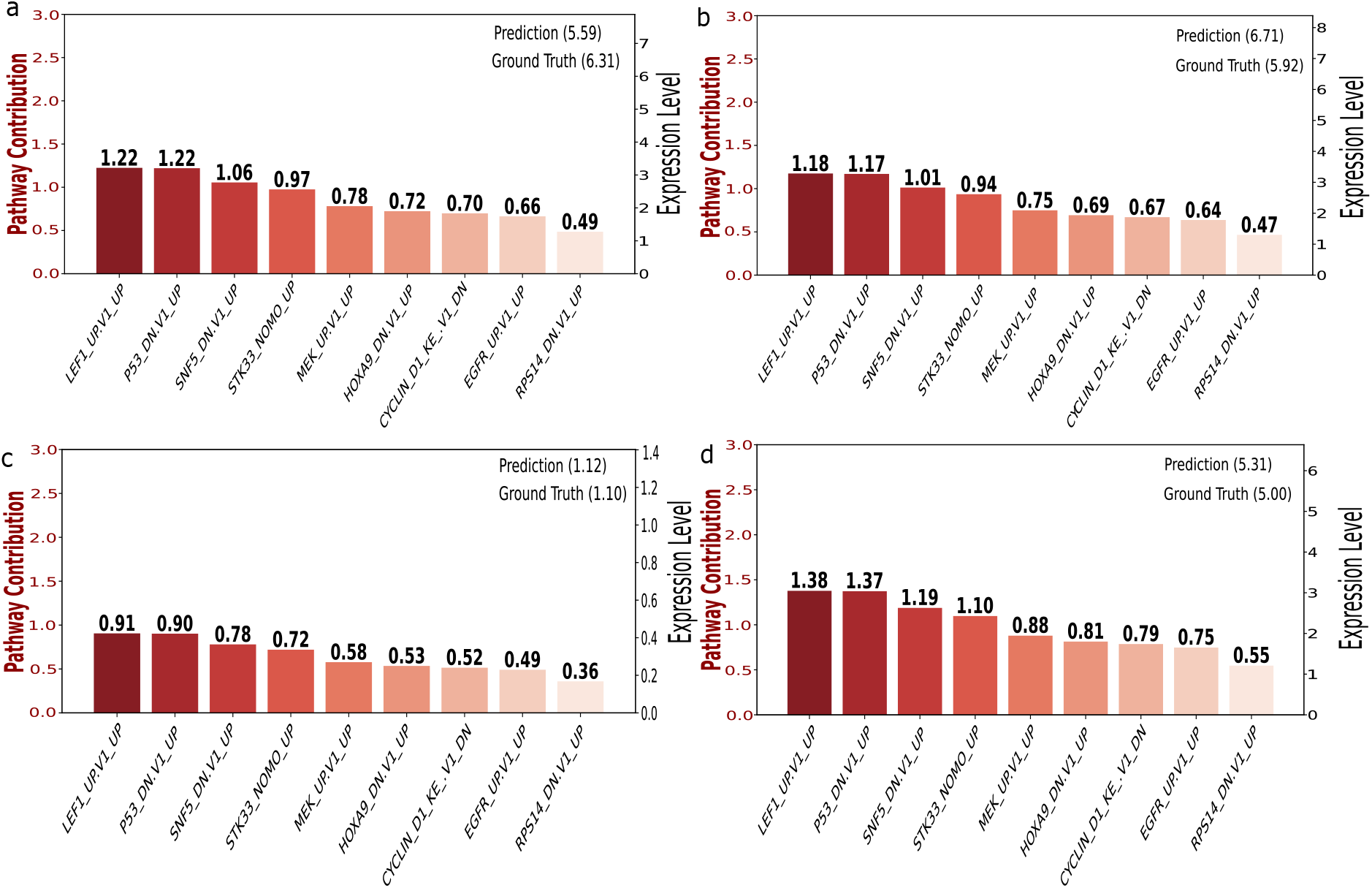
Interpretability analysis of gene predictions in the IDC dataset. Contribution analysis of four representative breast cancer biomarkers: **a** CENPF, **b** FOXA1, **c** MKI67 and **d** MLPH. EP-Booster accurately predicts these functionally diverse genes, with close agreement between predicted and true values. Notably, despite their distinct biological roles, a consistent core explanatory axis was observed across genes, with LEF1_UP.V1_UP and P53_DN.V1_UP ranked as the top contributors. This consistency suggests that the model captures stable histomorphological features rather than dataset-specific noise, reflecting suppression of Wnt and P53 signalling associated with a well-differentiated luminal cell state.

## Supplementary Tables

**Table S1.** Comparison with State-of-the-Art on the mean squared error (MSE) metric across IDC, SKCM, and PRAD datasets. Lower values indicate better performance. Results are averaged across cross-validation folds.

| Method | IDC (HVG) ↓ | SKCM (HVG) ↓ | PRAD (HMHVG) ↓ |
| --- | --- | --- | --- |
| ST-Net | 2.6918 | 3.9140 | — |
| HisToGene | 2.9903 | 2.7275 | 1.4619 |
| BLEEP | 2.6973 | 3.2473 | 2.4754 |
| TRIPLEX | — | — | 1.4819 |
| Stem | — | — | 1.4873 |
| Ours_R (ResNet50) | 2.5158 | 2.1104 | — |
| Ours_GP (ResNet50) | 2.4354 | 2.3830 | — |
| Ours_R (UNI) | 2.3575 | 1.5701 | 0.5442 |
| Ours_GP (UNI) | 2.3128 | 1.6127 | 0.5384 |
| Ours_R (CONCH) | — | — | 0.5599 |
| Ours_GP (CONCH) | — | — | 0.5472 |
| Ours_R (Virchow2) | 2.0904 | **1.3768** | **0.4894** |
| Ours_GP (Virchow2) | **2.0801** | 1.3895 | 0.4981 |

**Table S2.** Gene program lists for the SKCM and LUAD datasets derived from MSigDB pathways.

| SKCM | LUAD |
| --- | --- |
| P53 DN.V1 DOWN | HOXA9 DN.V1 UP |
| LEF1 UP.V1 UP | MEK UP.V1 UP |
| SNF5 DN.V1 UP | RPS14 DN.V1 UP |
| ESC V6.5 UP EARLY.V1 DOWN | EGFR UP.V1 UP |
| CAMP UP.V1 DN | ALK DN.V1 UP |
| RAF UP.V1 DOWN | AKT UP MTOR DN.V1 DOWN |
| P53 DN.V1 UP |  |
| NFE2L2.V2 |  |
| BMI1 DN MEL18 DN.V1 UP |  |
| BMI1 DN.V1 UP |  |
| STK33 SKM UP |  |
| GCNP SHH UP EARLY.V1 DOWN |  |

## Supplementary Text - Hyperparameters

We detail the hyperparameters used in our experiments, including learning rate, prototype dimensions, and prototype counts (for random initialization only).

### Learning Rate

Different learning rates are used based on the dataset and initialization strategy (*i*.*e*.,random and gene program-guided initialization):

- IDC (random): 0.01;
- LUAD (both): 0.001;
- IDC (gene program): 0.0001;
- SKCM (random, gene program): 0.0005;
- PRAD (random): 0.0001;
- PRAD (gene program): 0.00001.

### Prototype Dimension

The prototype dimension is set to 256 for most foundation models. Exceptions: ResNet-50 (512), CONCH and UNI (128).

### Number of Prototypes

The gene program-guided strategy automatically determines the number of prototypes. For random initialization:

- Most of the foundation models: 64;
- CONCH and UNI: 32.

## Supplementary Text - Pseudo Code

### Pseudo Code of EP-Booster

Algorithm 1 illustrates the training procedure of EP-Booster and how it generates the latent representation *Z*_Latent_ from image embeddings. During training, *Z*_Latent_ is obtained from H&E images via EP-Booster and used together with target gene expression to train a regression model. At test time, the trained EP-Booster generates *Z*_Latent_ from test H&E images, which is then fed into the trained regression model to predict gene expression.

#### Algorithm 1

A pipeline of EP-Booster.

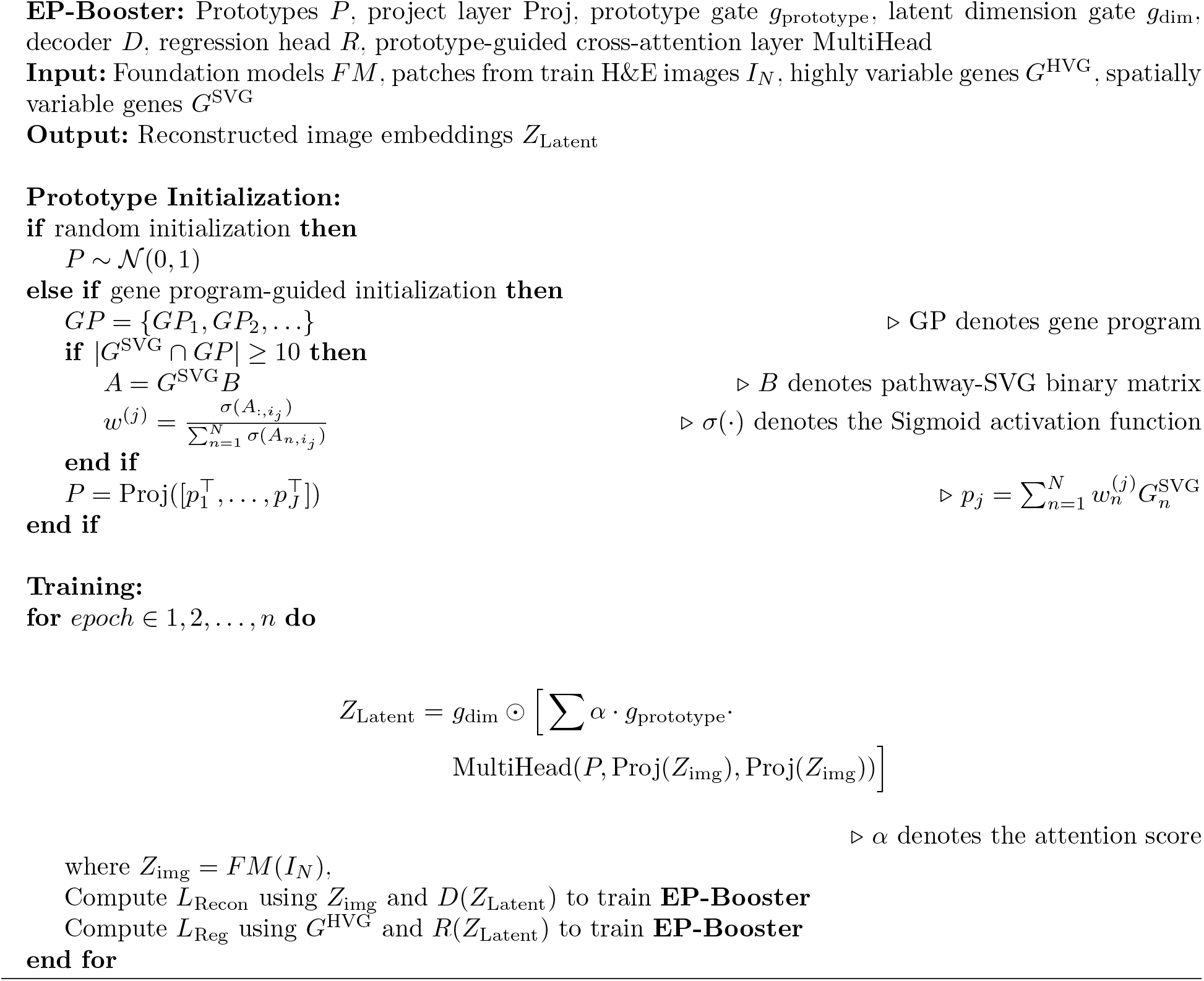

### Pseudo Code of Survival Analysis

#### Algorithm 2

Pipeline of survival analysis (TCGA evaluation)

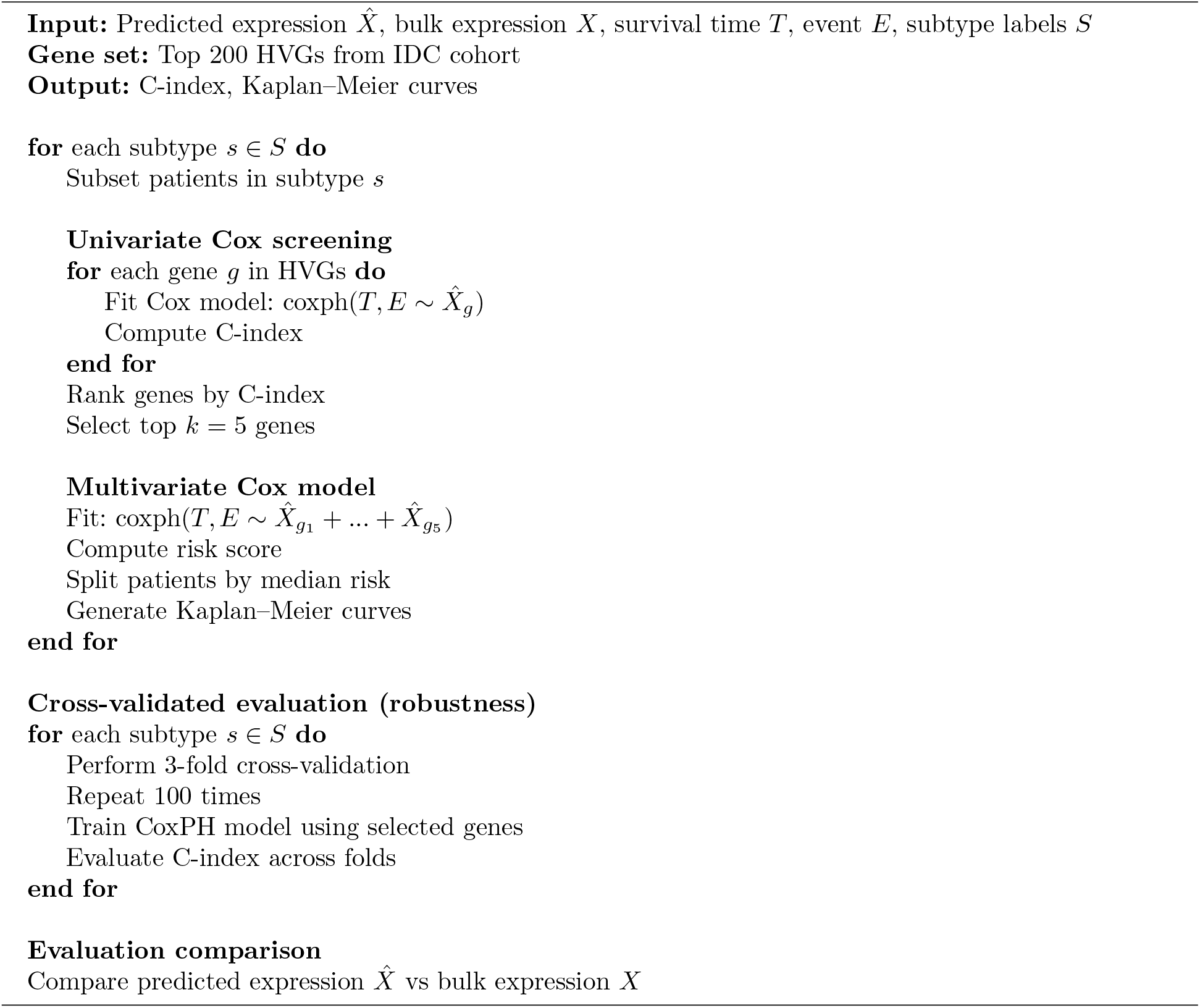

### Pseudo Code of Drug Response

#### Algorithm 3

Pipeline of drug response prediction (TransNEO evaluation)

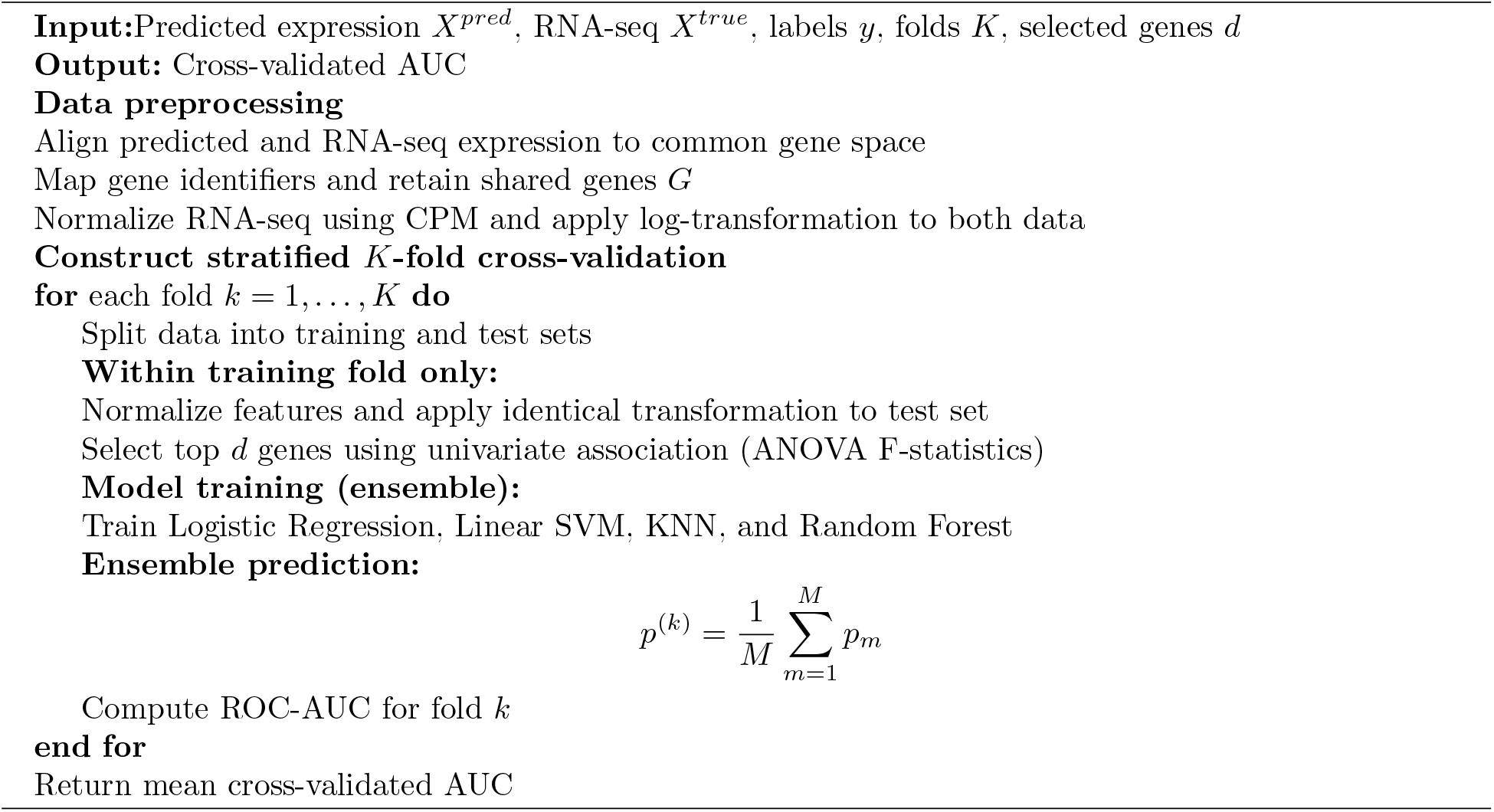

